# Genome-wide fitness profiling reveals flagellar rotation as an energetic liability during anaerobic maintenance in *Pseudomonas aeruginosa*

**DOI:** 10.64898/2026.09.04.749331

**Authors:** Kemal Demirer, Ryan A. Melnyk, Hans K. Carlson, Akihiro Okamoto, Adam M. Deutschbauer, Dianne K. Newman

**Affiliations:** Division of Biology and Biological Engineering, California Institute of Technology, Pasadena, California, USA; Division of Geological and Planetary Sciences, California Institute of Technology, Pasadena, California, USA; Research Center for Macromolecules and Biomaterials, National Institute for Materials Science, 1-1 Namiki, Tsukuba, Ibaraki, 305-0044, Japan; Faculty of Life and Environmental Sciences, University of Tsukuba, 1-1-1 Tennodai, Tsukuba, Ibaraki, 305-8577, Japan; Living Systems Materialogy (LiSM) Research Group, International Research Frontiers Initiative (IRFI), Tokyo Institute of Technology, Yokohama, Kanagawa, 226-8501, Japan; Environmental Genomics and Systems Biology Division, Lawrence Berkeley National Laboratory, Berkeley, CA; Department of Plant and Microbial Biology, University of California, Berkeley, CA

**Keywords:** *Pseudomonas aeruginosa*, phenazine, growth arrest, transposon sequencing, flagella, energy limitation

## Abstract

Bacteria in nature and disease frequently spend time in non-growing states, yet the genes that allow cells to survive without growing remain poorly understood. Using *Pseudomonas aeruginosa* strain PA14 as a model system to study non-growth powered by anaerobic phenazine cycling, we employed randomly barcoded transposon-insertion sequencing (RB-TnSeq) to identify the genes required for this maintenance state using a platform that sustains anaerobic survival via continuous phenazine reoxidation. We found 167 genes whose disruption altered survival, including genes involved in transcription, translation, protein quality control, and cell envelope maintenance. Comparing our results to published TnSeq data from other forms of energy-limited growth arrest (conditions where electron acceptor or carbon availability constrains energy conservation below that required for cell growth/division) in PA14 revealed that while a subset of genes are fitness determinants across distinct growth-arrested states, most are condition-specific. Notably, genes involved in flagellar regulation and assembly were broadly detrimental to survival under this maintenance condition. Leveraging a high-throughput electrochemical system that allows for quantitative and mechanistic dissection of the phenazine cycling-dependent maintenance state, we found that flagellar abundance influences cells’ survival, metabolic rate, and ATP levels. Moreover, removing the flagellar stator proteins MotAC, which are required for flagellar rotation but not assembly, reversed these defects, indicating that the energetic cost of flagella comes from their rotation rather than construction under these conditions. These results show that in a low-powered maintenance state, limiting energy-dissipating processes, such as proton-motive force loss through flagellar rotation, supports cell survival.

**Importance:** The genetic landscape governing non-growth survival under energy limitation, and the cellular processes that help or hinder it, remains largely uncharacterized. What does a cell need to do, and what must it avoid, to survive when growth-arrested? Answering this question is clinically important because non-growing bacteria are often antibiotic tolerant. Here, we used a genome-wide screen of *Pseudomonas aeruginosa* strain PA14, an opportunistic pathogen, to identify fitness determinants of viability during a non-growth state powered by anaerobic phenazine-cycling, a state mimicking the anoxic cores of biofilms. Our findings suggest that growth-arrested cells live at a bioenergetic knife’s edge, where an energy-dissipating process, like a rotating flagellum, can tip the balance between vitality and death, providing a possible bioenergetic rationale for the known repression of flagellar expression in mature biofilm cores.

## Introduction

It has long been appreciated that most microorganisms in natural environments exist in slow-growing or non-growing states (1, 2). *In situ*, microbes commonly experience nutrient limitation and other environmental stresses that restrict energy conservation and growth (3). Historically, however, studies of microbial physiology primarily have focused on oxic, nutrient-replete conditions that enable fast growth. While this disconnect is starting to be bridged (4–6), we still have a far more complete understanding of what underpins rapid growth compared to what sustains slow or non-growth. This knowledge gap is costly in the clinic: many antibiotics target active processes, allowing non-growing cells surviving with a low metabolic rate to tolerate these antibiotics (7–10). Understanding the genetic and physiological basis of non-growth physiology is thus of basic scientific interest and clinically important.

*P. aeruginosa* is an opportunistic pathogen that causes hundreds of thousands of deaths annually in patients with other lung comorbidities (11). A major contributor to its clinical persistence is its ability to form biofilms, multicellular aggregates in which cells experience oxidant limitation due to oxygen consumption outpacing its diffusion (12). In the anoxic cores of biofilms, cells enter a non-growth state, making them difficult to treat with conventional antibiotics (13–15). We have shown that *P. aeruginosa* PA14 can survive anaerobically in this microenvironment by utilizing phenazines—colorful, redox-active secondary metabolites—as terminal electron acceptors to support facilitated fermentation (16–18). Phenazines are reduced intracellularly and then transported outside the cell where they diffuse to and are reoxidized by distal oxidants. Returning to the cell in an oxidized form, they are taken up and re-reduced to repeat the cycle (16, 17, 19).

Though growth arrest has been studied in diverse bacterial pathogens, including *Legionella pneumophila* and *Salmonella enterica*, where the transition to non-growth is intimately linked to virulence and host adaptation (20, 21), arguably the best-studied case of bacterial survival during nutrient (*e.g.* carbon) limitation is that of *Escherichia coli* in stationary phase (5, 22, 23). In these studies, death of part of the population fuels growth of another, leading to a slow decay in net population viability. Though resource allocation shifts as growth slows (24, 25), it has been challenging to determine which cellular activities are abandoned in favor of others when growth stops entirely for want of a good system to study growth arrest quantitatively. Recently, we introduced a platform based on electron acceptor limitation utilizing phenazines that provides a tractable model for quantitative and mechanistic study of growth arrest (26).

Transposon-insertion sequencing (TnSeq) has emerged as a powerful approach for genome-wide fitness profiling under many conditions of interest, including non-growth states (27). Applied to *Pseudomonas aeruginosa* strain PA14 experiencing energy-limited growth arrest triggered either by carbon, nitrogen or oxygen limitation, TnSeq has identified both shared and condition-specific fitness determinants, revealing that regulatory genes are important during early survival while cell envelope-associated functions become beneficial later (28, 29). Advancements such as randomly-barcoded TnSeq (RB-TnSeq) have increased throughput significantly, enabling fitness profiling across hundreds of conditions in dozens of bacteria and revealing phenotypes for thousands of previously uncharacterized genes (30, 31). These studies have underscored that genetic determinants are highly context-dependent, such that the genes required for one condition (e.g. type of growth arrest) may not necessarily generalize to another, motivating genome-wide fitness profiling in different non-growth conditions.

Phenazine cycling-dependent anaerobic survival enables a maintenance state in which cells maintain viability without division. This state can be studied in the lab using high-throughput electrochemistry, where a constant oxidizing (0 mV vs. Ag/AgCl) potential is applied to anaerobically surviving cells to continuously reoxidize phenazines, and the current generated can be converted into a cellular power output. This latter feature of phenazine-dependent growth arrest permits quantitative assessment of the maintenance metabolic state, something difficult to achieve in other growth-arrested systems. Previous work from our lab has shown that cells in this state maintain a cell-specific metabolic rate ∼1000-fold lower than during aerobic growth in a nutrient-replete medium (26). Here, we report a genome-wide RB-TnSeq fitness screen of *P. aeruginosa* PA14 during anaerobic survival powered by phenazine-1-carboxamide (PCN)-cycling, revealing diverse physiological functions beneficial or harmful to this maintenance state and uncovering the energetic liability of flagellar rotation that can tip the balance between survival and death.

## Results

### An RB-TnSeq screen of anaerobic PCN-cycling survival identifies beneficial and harmful fitness determinants

To identify the genetic requirements for anaerobic survival by phenazine cycling, we performed a genome-wide fitness screen of *Pseudomonas aeruginosa* PA14 using an RB-TnSeq library constructed and introduced in this study (Supplemental Table 1). Cells from the library were grown aerobically in LB to early stationary phase (OD_500_ = 3.0) and inoculated into anoxic three-electrode bioreactors maintained under an oxidizing potential (0 mV vs. Ag/AgCl; see Methods) to continuously re-oxidize phenazine-1-carboxamide (PCN) reduced by the cells, and barcode abundances were tracked over seven days using RB-TnSeq (Methods) after aerobic outgrowth (2 hours in LB, ∼4 doublings) (Fig. 1).

**Fig. 1.**
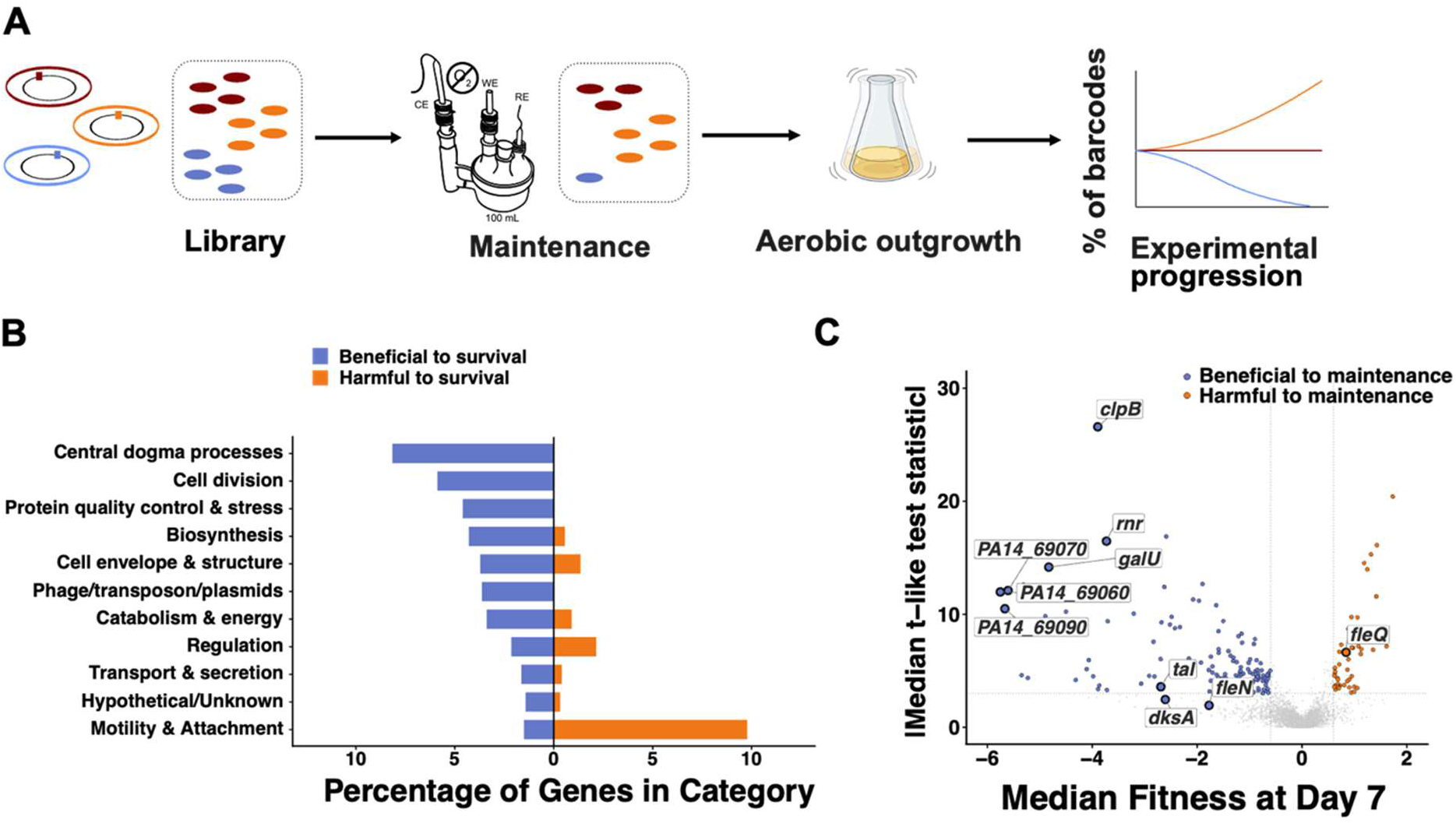
RB-TnSeq workflow and genome-wide fitness profiling of *P. aeruginosa* PA14 during anaerobic survival by phenazine cycling. **(A)** Schematic of the RB-TnSeq experimental workflow. A barcoded transposon insertion library was inoculated into anoxic bioreactors maintained under phenazine-1-carboxamide (PCN)-cycling conditions. At each sampling time point, cells were collected, aerobically outgrown in LB for ∼4 generations, and processed for barcode sequencing to calculate per-gene fitness relative to the initial inoculum. **(B)** Percentage of significantly beneficial (blue) and harmful (orange) genes within each functional category after 7 days of anaerobic survival under PCN-cycling conditions. Genes were classified as significant if the t-like test statistic exceeded 3 in absolute value in at least 2 of 3 replicates, with median fitness < −0.6 (beneficial) or > 0.6 (harmful). **(C)** Volcano plot showing median fitness scores at day 7 versus |median t-like test statistic| for all genes in the RB-TnSeq library. Blue points label genes beneficial to maintenance, orange points label those harmful to maintenance, and gray points label non-significant genes. Selected genes validated by independent assays are annotated. Dotted lines indicate fitness cutoffs (±0.6) and |t| = 3. Notably, *dksA* and *fleN* both met the fitness-magnitude but not the t-like test statistic significance threshold.

Fitness scores were calculated by comparing barcode abundances at day 7 to the day 0 sample, with genes scored as significantly beneficial (fitness < −0.6, |median t-like test statistic| > 3 in ≥2 of 3 replicates) or harmful (fitness > 0.6, |median t-like test statistic| > 3 in ≥2 of 3 replicates) to anaerobic maintenance. The fitness magnitude cutoff was adapted from Munro *et al.* (29), who used 0.58, which we rounded to 0.6, and the t-like test statistic was adapted from (30). The complete fitness dataset is provided in Supplemental File 1.

Because barcode abundances are measured after aerobic outgrowth, our experiment captures mutants that can survive anaerobically during phenazine-cycling as well as resume growth when cultured aerobically in LB. Mutants that survive the maintenance period but cannot re-enter aerobic growth thus appear as having fitness defects. Accordingly, we interpret fitness scores as reflecting requirements for the full maintenance-to-regrowth transition. Nevertheless, the phenotypes subsequently reported were confirmed for individual mutants in a subset of the genes identified by RB-TnSeq, independently of the liquid outgrowth step, using viability, metabolic rate, and/or ATP measurements. Resuscitation experiments in this system have shown that most CFU loss reflects cell death (26), so cultivability closely tracks viability under these conditions.

The screen identified 167 significant fitness determinants, of which 123 were beneficial and 44 were harmful to anaerobic PCN-cycling survival. We functionally annotated genes using the functional categories from the PA14 genome annotation (32), further grouping them into broad physiological categories for analysis (Fig. 1B; Supplemental File 1). To account for differences in category size across the genome, we depict enrichment as the percentage of genes within each functional category meeting significance thresholds, normalizing for differences in category representation in the PA14 genome.

Among the most strongly beneficial functional categories were central dogma processes, including transcription, RNA processing, and translation, as well as protein quality control & stress, a category encompassing chaperones and proteases that maintain protein homeostasis under stress (Fig. 1B). That genes spanning transcription, translation, and cell envelope maintenance are beneficial to survival highlights that anaerobic maintenance is not physiologically passive but requires investment in core cellular processes. Some of these beneficial genes may be the same as required during active growth, only regulated so they operate in an attenuated fashion; others may be distinct.

By contrast, the category with the highest proportion of genes harmful to survival was motility and attachment. Nearly 10% of genes in this category were significantly harmful to survival, and flagellar regulatory and structural genes were broadly enriched among these hits (Fig. 1C). Of the 44 genes harmful to survival, 16 are flagellar structural, assembly, or regulatory genes, representing the largest category among harmful genes. Notably, *fleQ*, the master transcriptional activator of flagellar biosynthesis (33, 34), and *fleN*, its anti-activator (35, 36), were harmful and beneficial to maintenance, respectively, suggesting that flagellar gene expression imposes a fitness cost during anaerobic maintenance. While *fleQ* met both fitness and significance criteria, *fleN* met the fitness-magnitude cutoff but not the t-like test statistic threshold in the primary screen. We nonetheless pursued it experimentally given its established role as the antiactivator of *fleQ*, allowing us to test both arms of the flagellar regulatory switch.

### Comparison of fitness determinants across growth-arrested conditions reveals that PCN-cycling survival is a physiologically distinct state

Having identified diverse fitness determinants of PCN-cycling survival, we next asked whether these requirements are specific to this maintenance state or generalizable to other forms of growth arrest. We compared our dataset to previous TnSeq fitness data from our lab where we studied *P. aeruginosa* PA14 under oxygen-limited (O_2_-lim) and carbon-limited (C-lim) growth arrest (28). Datasets were merged on PA14 locus tags, and genes were classified as beneficial to maintenance using a consistent threshold (fitness < −0.6, p < 0.05, limma moderated t-test) across all three conditions. Because this analysis evaluates replicate-level variance, it may identify genes with consistent effects across replicates that do not meet the per-insertion consistency threshold used in the PCN-only screen (Fig. 1).

Most fitness determinants were condition-specific, with 194 genes uniquely required under PCN-cycling conditions compared to 123 unique to O_2_-lim and 89 unique to C-lim (Fig. 2A). This finding suggests that different forms of maintenance are not physiologically equivalent: each condition imposes different constraints, and the genetic requirements for survival reflect those differences rather than a universal maintenance program, consistent with prior work demonstrating that fitness determinants are highly context-dependent (28). We hasten to note, however, that the medium used in our experiments differed slightly from that used in the earlier growth-arrest TnSeq study (*e.g.* glucose vs. pyruvate as the carbon source), as did the TnSeq library, so some of the variation between the datasets may also reflect these differences. Additionally, PCN itself may exert toxicity effects that influence fitness scores independent of maintenance physiology.

**Fig. 2.**
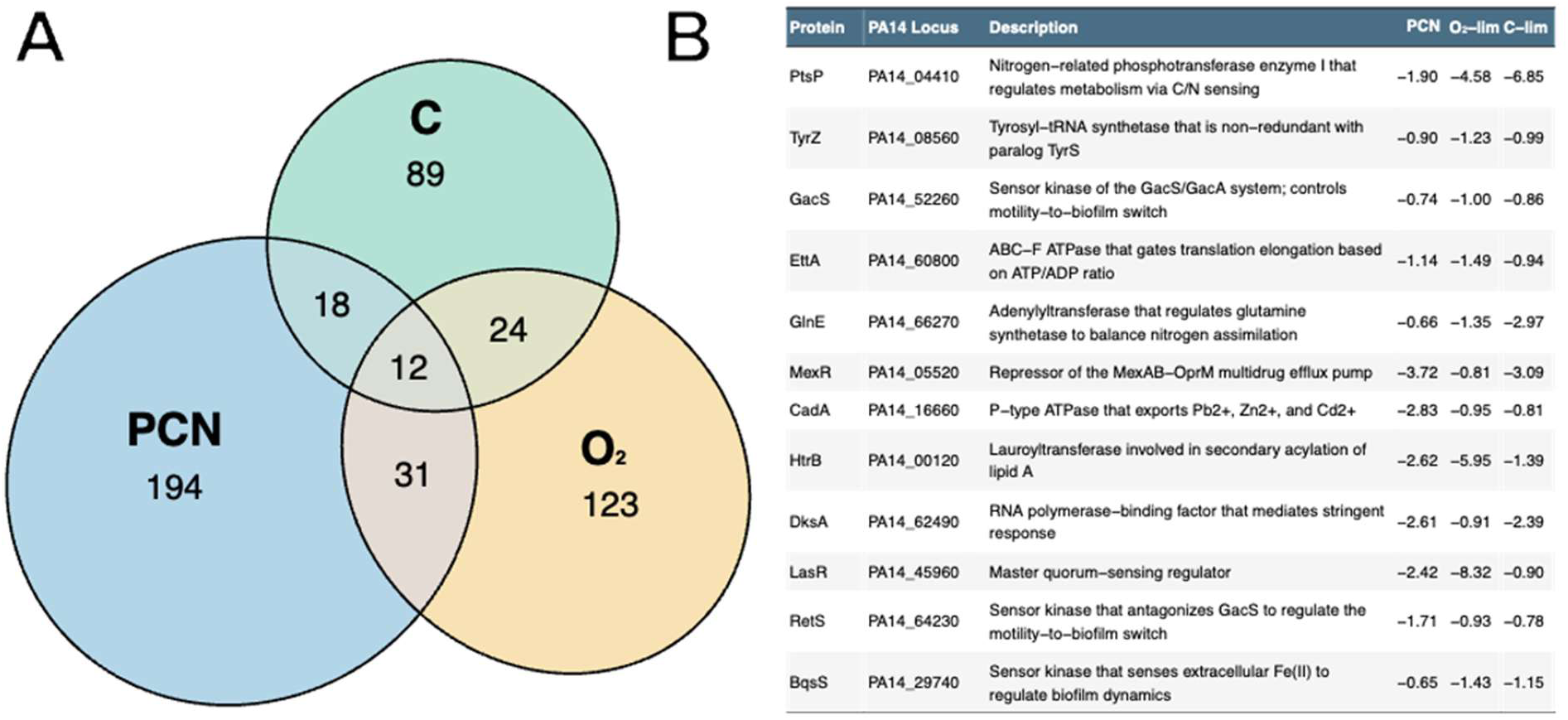
PCN-cycling survival requires a largely distinct set of fitness determinants compared to other growth-arrest conditions. **(A)** Euler diagram showing the overlap of determinants beneficial to maintenance (fitness < −0.6, p < 0.05) identified under PCN-cycling (this study), oxygen-limited (O_2_-lim), and carbon-limited (C-lim) conditions. O_2_-lim and C-lim fitness data are from Basta *et al.* (28). Numbers indicate genes meeting significance thresholds uniquely or jointly in each condition. **(B)** Table of the twelve genes significant across all three conditions, showing PA14 locus tags, functional descriptions, and fitness scores under each condition. Fitness scores are shown as log_2_ values.

Nevertheless, 31 genes were shared between PCN-cycling and O_2_-lim conditions, which we interpret as a strong signal pointing to common processes. Overlaps of 24 and 18 genes were observed between C-lim and the O_2_-lim or PCN-cycling conditions, respectively (Fig. 2A). The ABC transporter operon (PA14_69060, PA14_69070, PA14_69090) was among the strongest fitness determinants in both PCN-cycling and O_2_-lim datasets (Supplemental File 2). Notably, this operon was also transcriptionally upregulated during malonate utilization in a third independent study (37), a carbon source that induces anaerobic stress responses in PA14.

Twelve genes met significance thresholds across all three conditions, representing a minimum candidate core set of genes required to survive diverse forms of growth arrest (Fig. 2B, Supplemental File 2). All twelve exhibited negative fitness scores across conditions. Notably, the core set is dominated by sensory and regulatory genes rather than metabolic enzymes. *ptsP*, with its largest defect under C-lim (-6.85), integrates carbon and nitrogen status to regulate metabolism (38), while *glnE* tunes glutamine synthetase activity to balance nitrogen assimilation (39). The GacS/GacA two-component system, which controls the motile-to-biofilm lifestyle switch and virulence, is represented by both of its opposing sensor kinases, *gacS* and *retS* (40). Additional regulatory functions include *bqsS*, which senses extracellular Fe(II) (41, 42), *lasR*, the master quorum-sensing regulator (43), and *dksA*, which mediates the stringent response through direct interaction with RNA polymerase (44). Beyond regulation, *ettA* gates ribosomal entry into translation elongation in proportion to the cellular ATP/ADP ratio (45, 46), and *htrB*, a lauroyltransferase involved in lipid A acylation, supports outer membrane integrity (47, 48). The remaining core genes, *tyrZ*, *cadA*, and *mexR*, are also included in Fig. 2B.

### Validation of RB-TnSeq fitness determinants confirms diverse physiological requirements for anaerobic survival

To validate the findings of our screen, we selected a subset of top hits spanning diverse functional categories (Fig. 1C), prioritizing genes for which clean deletion or transposon insertion mutants were immediately available, and assessed their survival in our high-throughput electrochemical system (26). These included *rnr*, encoding a processive 3’→5’ exoribonuclease involved in RNA degradation and quality control; three subunits of the aforementioned ABC transporter of unknown function (*PA14_69060*, *PA14_69070*, *PA14_69090*); *tal*, encoding transaldolase; *clpB*, encoding an AAA+ disaggregase chaperone; *dksA*, encoding an RNA polymerase-binding transcriptional regulator; and *galU*, encoding UTP-glucose-1-phosphate uridylyltransferase (Fig. 3B). Although *dksA* did not meet the t-like test statistic threshold in the primary PCN-cycling screen, we selected it for validation based on its identification as a fitness determinant shared in our comparative analysis (Fig. 2B) and because it passed the fitness threshold.

**Fig. 3.**
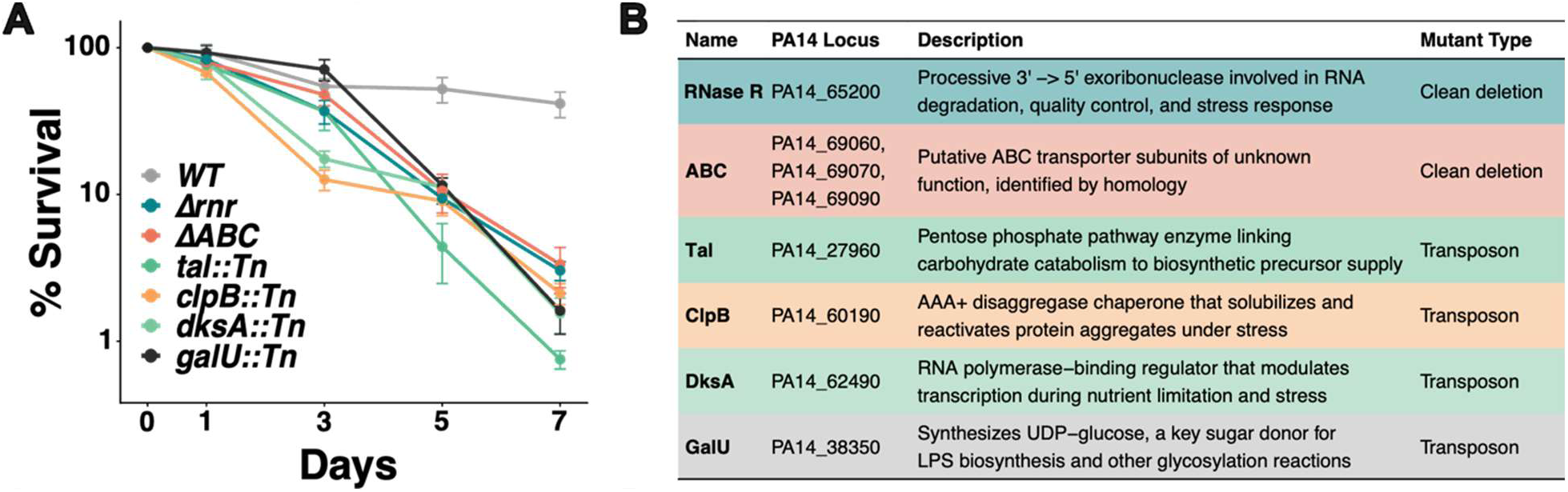
Transposon insertion and clean deletion mutants in genes spanning diverse functional categories show survival defects during anaerobic PCN-cycling survival. **(A)** Survival of WT and the indicated mutant strains over seven days of anaerobic PCN-cycling survival, expressed as percent survival relative to day 0. Data points represent the mean of n = 6 biological replicates; error bars indicate standard error of the mean. **(B)** Table summarizing the validated genes, their PA14 locus tags, functional descriptions, functional categories, and mutant types used for validation.

We assessed the survival of clean deletion or transposon insertion mutants in each gene (clean-deletion mutants in ABC transporter (28) and *rnr*, Tn mutants from the Ausubel PA14 non-redundant mutant library (49) in *tal*, *clpB*, *galU*, and *dksA*) over seven days under anoxic PCN-cycling conditions, along with WT PA14 (Fig. 3A). Transposon mutants were PCR-verified using primers flanking the insertion region. All six mutants exhibited reduced survival relative to WT, confirming their importance for anaerobic maintenance. The severity of survival defects varied across mutants: Δ*rnr* cells showed the most severe loss of viability by day 7, while *galU*, *clpB*, *dksA*, and *tal* mutants displayed intermediate defects, and disruption of the ABC transporter resulted in a more modest but reproducible reduction in viability. Notably, *dksA* and *clpB* mutants lost viability more rapidly than the other four validated hits, which declined more gradually over the seven-day time course.

### Flagella are a burdensome energetic drain during anaerobic maintenance

An unexpected finding of our screen was the harmful effect of flagellar genes, motivating us to investigate the energetic consequences of flagellar abundance during anaerobic PCN-cycling survival. Spontaneous non-motile mutants, including GacS/GacA mutants that lose motility through disruption of the motile-to-sessile switch (40), are commonly observed in *Pseudomonas* in both laboratory and field settings (50), providing context for the energetic cost of flagellar rotation we observe under maintenance conditions. Furthermore, because motility is regulated by a bistable switch, some cells in the inoculum will be motile and others non-motile upon transition into stationary phase. Non-motile cells will be protected from the energetic cost of flagellar rotation, while motile cells will suffer this cost as they enter the maintenance state. The master flagellar activator, FleQ, as well as other flagellar structural/regulatory genes, is harmful to survival under this condition while FleN, the flagellar anti-activator in *P. aeruginosa*, is beneficial.

To investigate the effects of these genes on maintenance metabolism, we made clean deletions of these regulators in the Δ*phz* genetic background, where PA14 cells cannot make phenazines. For survival experiments using these strains, all phenazines were provided exogenously to enable consistency in phenazine concentration across experiments. We measured survival, cell-specific metabolic rate (CSMR), and per-cell ATP concentrations in Δ*fleQ*Δ*phz* (flg-null) and Δ*fleN*Δ*phz* (flg-OP, overproducing) mutants relative to the Δ*phz* parent strain to test the hypothesis that flagellar abundance impacts the energy economy of maintaining cells.

Flg-null cells survived significantly better than Δ*phz* over seven days, while flg-OP cells showed survival defects compared to the parent strain (Fig. 4A). These phenotypes were complemented by chromosomal reintegration of *fleQ* (flg-null +) and *fleN* (flg-OP +) at the attTn7 site, confirming that the observed effects were due to loss of each regulator (Fig. 4A). Consistent with an energetic cost associated with flagellar abundance, flg-null cells exhibited a significantly lower CSMR than Δ*phz*, while flg-OP cells showed a significantly elevated CSMR (Fig. 4B). Per-cell ATP concentrations mirrored these findings: flg-null cells maintained significantly higher ATP levels than Δ*phz* on day 1, whereas flg-OP cells had significantly lower ATP concentrations. Complementation restored both CSMR and ATP levels to those of the parent strain in both backgrounds (Fig. 4C). Interestingly, differences in per-cell ATP levels were observed on day 1 of the experiment, before survival differences had manifested.

**Fig. 4.**
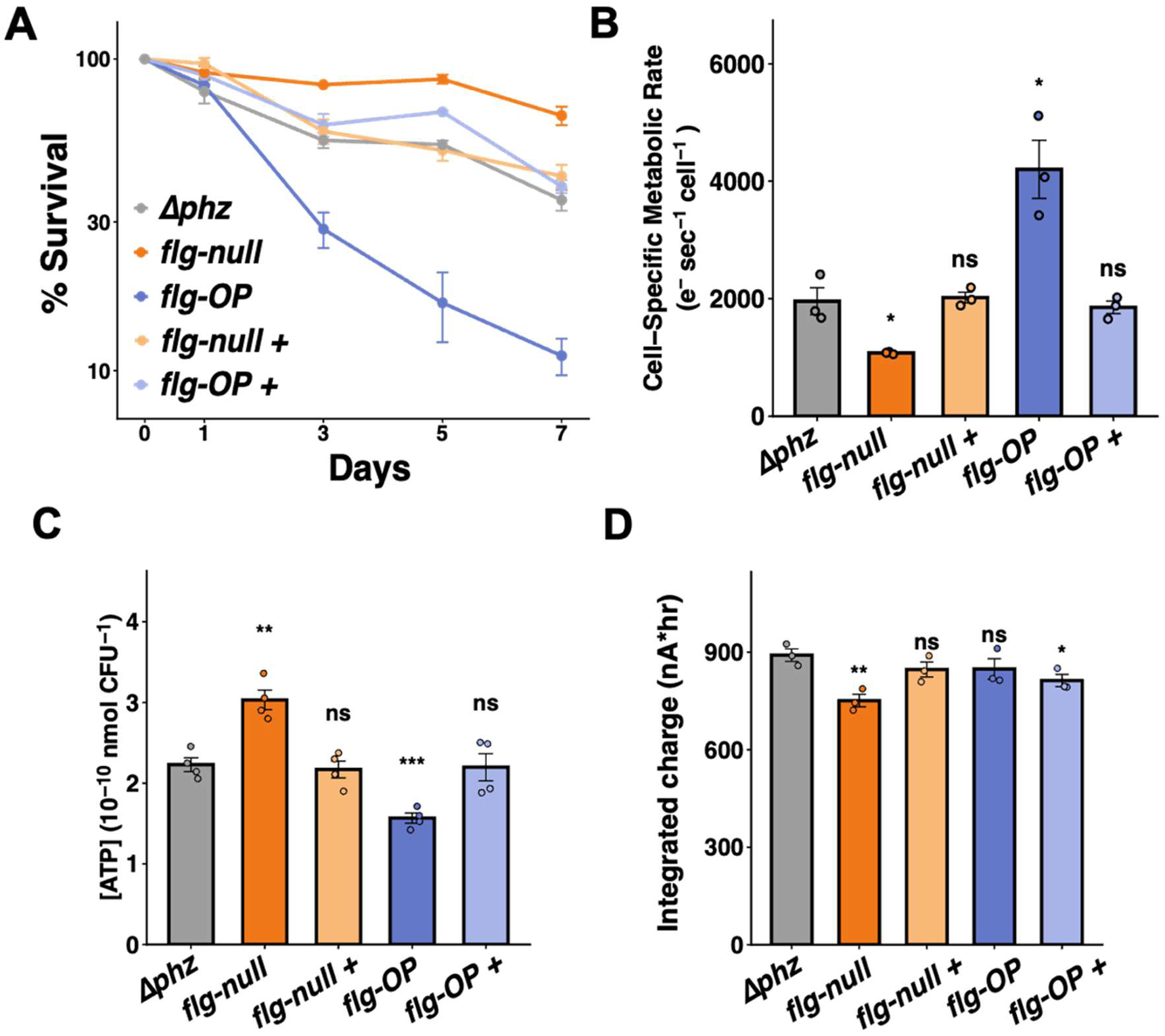
Flagellar abundance tunes cell-specific metabolic rate and ATP levels during anaerobic survival by phenazine cycling. **(A)** Survival of *Δphz*, flg-null, flg-null +, flg-OP, and flg-OP + strains over seven days of anaerobic PCN-cycling survival, expressed as percent survival relative to day 0. Data points represent the mean of n = 3 biological replicates; error bars indicate standard error of the mean. **(B)** Cell-specific metabolic rate (CSMR, electrons·cell⁻¹·sec⁻¹) of the indicated strains measured during anaerobic PCN-cycling survival. **(C)** Per-cell ATP concentrations (nmol CFU⁻¹) of the indicated strains on day 1 of anaerobic survival. **(D)** Integrated charge (nA·hr) over the first 24 h of anaerobic survival. For panels B and D bars represent the mean of n = 3 biological replicates while bars in panel C represent the mean of n = 4 biological replicates; error bars indicate SEM; individual data points are shown. Statistical comparisons are relative to *Δphz*; *, p < 0.05; **, p < 0.01; ***, p < 0.001; ns, not significant (Student’s t-test).

To rule out the possibility that these ATP differences reflect strain-specific differences in PCN redox cycling rather than flagellar ATP drain, we compared the integrated charge over the first 24 hours across strains, the same window after which ATP was measured and in which CFUs remained comparable across strains (Fig. 4D). Integrated charge was not significantly different between flg-OP and the *Δphz* parent, despite the significantly lower per-cell ATP in flg-OP cells, indicating that the ATP deficit does not arise from reduced electron flux through the extracellular electron transfer (EET) pathway. flg-null cells showed a significant reduction in integrated charge relative to *Δphz*, consistent with their lower cell-specific metabolic rate; however, per-cell ATP was elevated, not reduced, in flg-null cells, which indicates that EET rate is not the primary determinant of the observed ATP differences. These results indicate that the per-cell ATP differences reflect differential ATP consumption by flagella rather than differences in ATP generation via different rates of EET.

Taken together, these data demonstrate that flagellar abundance is a key determinant of the energy economy of maintaining cells. These observations raised the question: does the energetic burden of flagella arise primarily from the cost of biosynthesis, or from dissipation of the proton motive force during flagellar operation?

### Flagellar rotation, not biosynthesis, imposes an energetic burden during anaerobic maintenance

To distinguish whether the energetic burden of flagella during anaerobic PCN-cycling survival arises from costs associated with flagellar biosynthesis or from the dissipation of the proton-motive force (PMF) during flagellar rotation, we constructed clean deletions of *motA* and *motC* in the *Δphz* and flg-OP backgrounds. MotA and MotC are stator subunits of the flagellar motor that couple PMF dissipation to torque generation; cells lacking these subunits assemble flagella but cannot rotate them, allowing us to distinguish the cost of rotation from that of biosynthesis (51, 52). As a control, *motAC* deletion alone in the *Δphz* background improved survival, reduced CSMR, and elevated ATP relative to the *Δphz* parent strain, suggesting that the primary energetic burden of flagella during maintenance arises from their rotation rather than their biosynthesis.

We then asked whether the same logic explains the survival defect of flg-OP cells. Deletion of *motAC* in the flg-OP background fully rescued the survival, CSMR, and ATP defects of flg-OP cells, restoring all three parameters to levels comparable to flg-null cells that do not express flagellar genes (Fig. 5A-C). Integrated charge over the first 24 hours was not significantly different among flg-null, flg-OP, and flg-OP *ΔmotAC* strains (Fig. 5D), confirming that the ATP differences between these strains do not reflect differences in EET rate. Critically, deletion of *motAC* restored per-cell ATP to flg-null levels without altering integrated charge, directly demonstrating that flagellar rotation drains ATP per cell rather than reducing the rate of ATP generation via PCN cycling. *ΔmotACΔphz* cells also resembled flg-null cells in survival, CSMR, and ATP levels (Supp. Fig. 1A–C), indicating that the cost of flagella during maintenance is attributable to flagellar rotation rather than presence, regardless of flagellar abundance. This rescue suggests that flagella are already made prior to entrance into the maintenance state, and that the energetic liability of flagella during maintenance arises from PMF dissipation during rotation rather than the cost of flagellar biosynthesis.

**Fig. 5.**
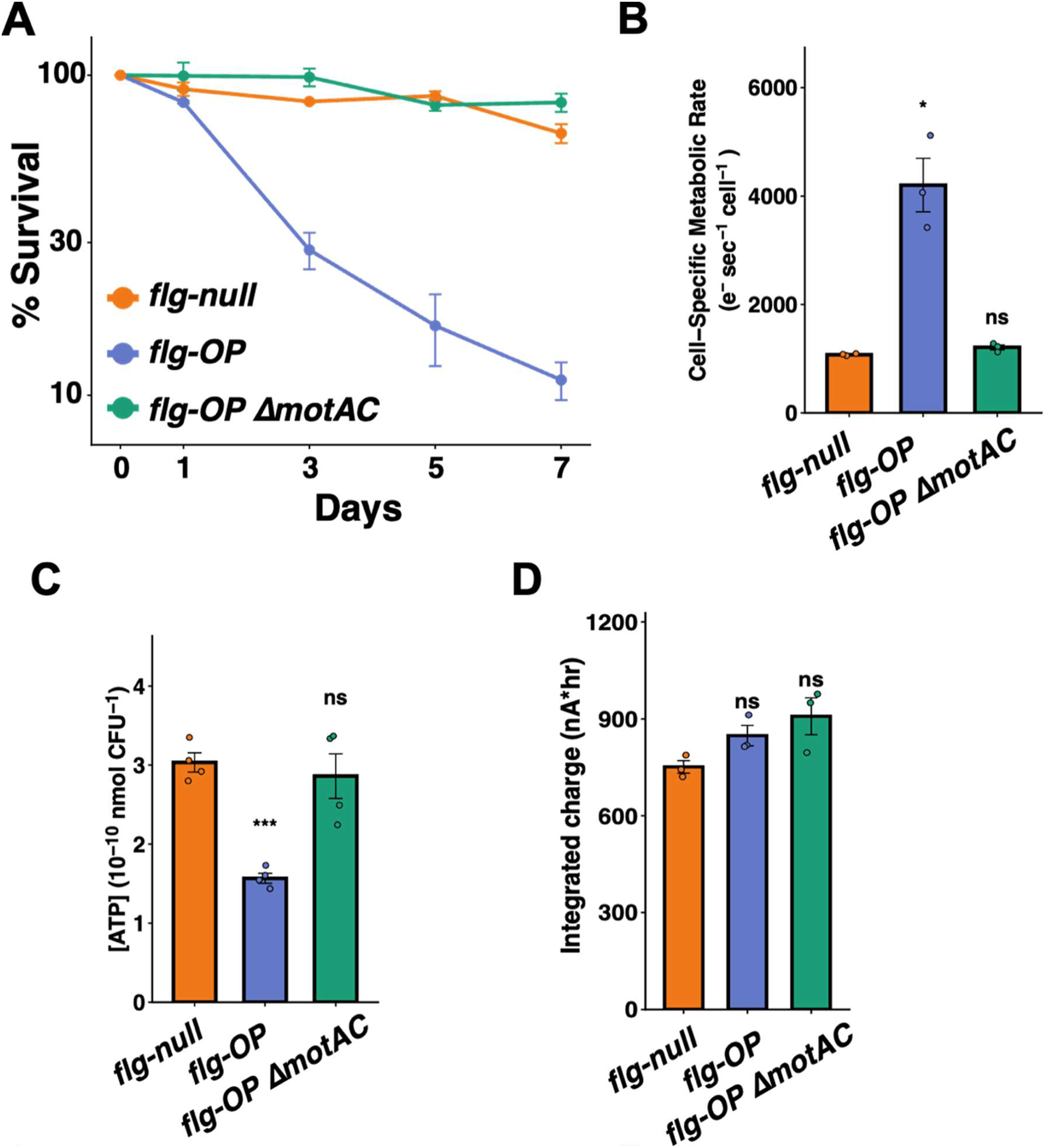
Loss of *motAC* restores cell-specific metabolic rate and ATP levels in a *ΔfleN* background during maintenance. **(A)** Survival of flg-null, flg-OP, and flg-OP *ΔmotAC* strains over seven days of anaerobic PCN-cycling survival, expressed as percent survival relative to day 0. Data points represent the mean of n = 3 biological replicates; error bars indicate standard error of the mean. **(B)** Cell-specific metabolic rate (CSMR, electrons·cell⁻¹·sec⁻¹) of the indicated strains measured during anaerobic PCN-cycling survival. **(C)** Per-cell ATP concentrations (nmol CFU⁻¹) of the indicated strains on day 1 of anaerobic survival. **(D)** Integrated charge (nA·hr) over the first 24 h of anaerobic survival. For panels B and D bars represent the mean of n = 3 biological replicates while bars in panel C represent the mean of n = 4 biological replicates; error bars indicate SEM; individual data points are shown. Statistical comparisons are relative to *ΔfleQΔphz*; *, p < 0.05; **, p < 0.01; ***, p < 0.001; ns, not significant (Student’s t-test).

## Discussion

Survival under energy-limited conditions requires not only that cells have access to sufficient energy to maintain viability but that they manage their energetic expenditures prudently. Using a genome-wide RB-TnSeq fitness screen of *P. aeruginosa* PA14 surviving by anaerobic phenazine-cycling, we identified fitness determinants spanning diverse physiological functions, including core cellular processes such as transcription, translation, protein quality control, and cell envelope maintenance. Among the harmful hits, flagellar regulatory and structural genes stood out, with the master activator *fleQ* among the most harmful and its anti-activator *fleN* among the most beneficial. This pattern prompted us to investigate the energetic consequences of flagellar abundance during maintenance. We found that flagellar rotation impedes survival by dissipating the PMF, and that its abrogation is sufficient to rescue survival, metabolic rate, and ATP levels. When cells exist close to their bioenergetic limit, the energetic burden of a single constitutive process thus appears to be able to tip the balance between life and death.

Our comparative analysis of fitness determinants for growth arrest across three different conditions (C-lim, O_2_-lim and PCN-cycling) reveals both shared and distinct features. For example, the ABC transporter operon identified here was significantly beneficial to survival under the two oxidant-limited conditions but not under carbon limitation. A broadly conserved core of twelve genes was required for survival across all three conditions (Fig. 2B). Among these, *ettA*, which gates ribosome entry into the translation elongation cycle through an interaction dependent on the cellular ATP/ADP ratio (45, 46), is particularly interesting: its recurrence across all three conditions suggests that ATP-dependent translational regulation may be a general strategy for managing energy economy during growth arrest. *ptsP*, the sensor enzyme of the nitrogen-related phosphotransferase system (PTS^Ntr^) that integrates carbon and nitrogen status to regulate diverse aspects of metabolism and stress response (53), implicates metabolic sensing as a broadly conserved requirement across growth-arrested contexts, a conclusion reinforced by Munro *et al.*, who identified *ptsP* and *ptsO* as among the strongest fitness determinants across carbon- and nitrogen-limited starvation in *P. aeruginosa* (29). This pattern extends to *glnE*, which adenylylates glutamine synthetase to tune nitrogen assimilation in response to cellular nitrogen status (39), further implicating nitrogen homeostasis as a requirement across distinct growth-arrested conditions. The minimal core set of genes also includes regulators of lifestyle switching and environmental sensing. Both *gacS* and *retS*, the opposing sensor kinases of the GacS/GacA two-component system that controls the motile-to-biofilm switch and virulence (40), were required across all three conditions. The identification of both kinases as core fitness determinants validates that the anaerobic phenazine-cycling system faithfully captures key features of the biofilm lifestyle, and is further supported by the recurrence of *lasR*, the master quorum-sensing regulator (43), and *bqsS*, which senses extracellular Fe(II) and has been implicated in regulating biofilm dynamics (41, 42). Additionally, *dksA*, which binds RNA polymerase to mediate the stringent response (44), points to transcriptional repression of growth-related processes as a broadly conserved survival strategy during growth arrest.

Consistent with our prior research demonstrating cells in the PCN-cycling condition are operating at their basal metabolic power (26), the opposing phenotypes of flg-null and flg-OP cells imply that cells in the maintenance state are at a bioenergetic knife’s edge. In this state, small changes in energy expenditures can have large consequences for survival. Under PCN-cycling conditions, the PMF is mainly generated by hydrolyzing ATP at the F_0_F_1_-ATPase, and cells are sensitive to drugs that disrupt the PMF (17). Here, we show that changes in PMF dissipation are correlated with changes in survival outcomes as well as cellular ATP levels. While ATP thresholds for viability loss have been observed in other energy-limited systems (28, 54, 55), our electrochemical approach enables measurement of the cellular power output at which this threshold occurs. Previously, we found this threshold is 3-4 orders of magnitude below that of fast growth (26), and here we show that a single energy-dissipating process, flagellar rotation, can push cells past it. A sensitivity to dysregulation of cellular processes that exact an energetic toll may also explain our observation that *dksA* and *clpB* mutants lost viability more rapidly than other mutants (Fig. 3A). DksA loss would prevent transcriptional adaptation at the start of the stringent response (44), and ClpB loss would permit protein aggregates to accumulate (56); both of these phenomena would present challenges to the cell that would require sufficient ATP to solve.

The CSMR data directly quantify the bioenergetic cost of flagellar rotation under PCN cycling: the difference between the *Δphz* strain (1,958 electrons·cell⁻¹·s⁻¹) and the non-rotating *ΔmotACΔphz* strain (1,309 electrons·cell⁻¹·s⁻¹) yields a rotation cost of 649 electrons·cell⁻¹·s⁻¹ – 33% of the total energy budget. Flagellar rotation is inherently expensive; Schavemaker and Lynch (57) estimated that bacterial flagellar operating costs are on the same order as maintenance energy requirements. Under aerobic growth, the cell’s energy budget substantially exceeds maintenance needs, accommodating flagellar costs alongside other growth costs. Under PCN-cycling maintenance, where metabolic power is low, spending a third of the budget reduces the energy available for maintenance. In biofilm cores, where oxygen is depleted and PCN cycling is the sole energy-conserving mechanism, this favors non-motility as a fitness strategy.

That flagellar activity can become an energetic liability under energy stress has been suggested by recent work by others. Leitner *et al.* showed that *ΔmlaE V. cholerae* cells, which maintain ∼50% lower cellular ATP than wild type, lose cultivability in stationary phase, a defect rescued by spontaneously arising suppressor mutations that inactivate flagellar biosynthesis and partially restore cellular ATP (58). Munro *et al.* similarly found that loss of flagellar motility in carbon- and nitrogen-starved *P. aeruginosa* frees resources for other biosynthetic activities, and that *fleN* loss imposes one of the strongest fitness defects observed across starvation conditions in this organism, consistent with our finding that *fleN* is beneficial under PCN-cycling conditions (29). Most recently, Migueles-Lozano *et al.* showed that hyperflagellation due to *ΔfleN* reduces growth, virulence, and competitive fitness in *P. aeruginosa*, further demonstrating that the FleQ-FleN circuit actively balances flagellar number against fitness costs (59). Our *ΔmotAC* result builds on these observations by dissecting the source of the cost: it is specifically PMF dissipation during rotation, not the biosynthetic investment in building the flagellum, that dictates the survival outcome at the bioenergetic margin.

The temporal ordering we observe here—bioenergetic changes preceding survival changes (*e.g.* ATP differences arising before viability changes during phenazine-supported anaerobic survival (17)) —parallels NADH/NAD^+^ dynamics in *P. aeruginosa* colony biofilms, where intracellular NADH/NAD^+^ ratios peak just before the induction of colony wrinkling, a morphological adaptation that maximizes oxygen access in the energy-limited biofilm interior (60). These patterns make sense given that phenazines facilitate NADH reoxidation when cells are oxidant-limited, which in turn facilitates flux through ATP-generating pathways (17, 61). The fitness consequences of flagellar regulation identified are relevant for non-growing cells in biofilm cores. Flagella are broadly downregulated during biofilm formation in *P. aeruginosa* (62) as well as in the lungs of individuals living with cystic fibrosis (63), a process driven in part by c-di-GMP-mediated repression of flagellar gene expression through FleQ (64). Our data suggest that this downregulation may enable bioenergetic prudence (i.e. PMF preservation). In addition, flagellar loss may confer an additional advantage in the CF lung: flagellin is a potent TLR5 agonist (65), and flagellum-deficient *P. aeruginosa* shows enhanced virulence in CF infection models, whereas non-motile but flagellated mutants do not (66). This suggests that *in vivo*, complete flagellar loss, not solely loss of rotation, may be optimal, providing both bioenergetic and immune evasion benefits.

Taken together, our findings indicate that energy-limited survival is exquisitely sensitive to the costs of individual cellular processes (3, 57). While flagellar rotation serves as a proof-of-concept, the broader fitness landscape we describe suggests that other processes may follow similar logic. Going forward, identifying the factors that permit cells to prioritize survival over growth represents an exciting cell biological frontier. The quantitative framework developed here can be applied to other energy-dissipating processes, and future work measuring how cells manage essential yet expensive energetic processes under different growth-arrested conditions will help us develop a quantitative bioenergetic understanding of bacterial maintenance.

## Materials and Methods

### Bacterial strains and culturing conditions

The strains and plasmids used in this study can be found in Supplemental Table 1. For strain maintenance and routine growth, *E. coli* and *P. aeruginosa* PA14 were grown at 37°C in lysogeny broth (LB) (Difco) containing 5 g/L yeast extract, 10 g/L tryptone, 10 g/L NaCl, and 15 g/L agar for culturing on solid medium. During aerobic growth, cultures were shaken at 250 rpm on a standard shaker (VWR). For anaerobic culturing of *P. aeruginosa*, 20 mM KNO_3_ was added to the LB medium. Single transposon mutants retrieved from the non-redundant transposon library (49) were verified by colony PCR using primers that flank the insertion site.

### Mutant construction and complementation in *P. aeruginosa*

Clean deletion mutants in *P. aeruginosa* were constructed according to the allelic exchange method described in (67) using ∼1 kb regions flanking the gene(s) of interest in the pMQ30 plasmid. Plasmids were constructed using standard Gibson assembly reactions (NEB). All primers used in this work were synthesized by Integrated DNA Technologies. Complement strains were constructed by introducing the gene *in trans* at the *att*Tn7 site downstream of *glmS* using the pJM220 plasmid. Complementation strains were designed to have each gene expressed by its native promoter in the genome (150 bp upstream of the 5’ end of the coding sequence). Transposon mutants were drawn from the PA14 non-redundant transposon library (49). All primers and plasmids used in this study can be found in Supplemental Table 1.

### RB-TnSeq in Pseudomonas aeruginosa PA14

To apply RB-TnSeq in *P. aeruginosa* PA14, we generated the barcoded *mariner* transposon delivery vector pAD280_NN1 containing millions of DNA barcodes and a gentamicin resistance cassette. pAD280_NN1 is derived from the transposon delivery vector pBT20 (68), which has been used previously for transposon mutagenesis in *Pseudomonas aeruginosa* (69–71). To make pAD280_NN1, we first PCR-amplified around the pBT20 plasmid with oAD1664 and oAD1665, followed by a Gibson assembly reaction to circularize the PCR product. The resulting plasmid (pAD280) is identical to pBT20 except it contains two BbsI sites for incorporating the DNA barcodes into the transposon via Golden Gate assembly, using a previously described strategy (72). To generate the PCR product with the random DNA barcodes, we PCR amplified ofeba282 with oAD1657 and oAD1658. A Golden Gate reaction with this PCR product and pAD280 was used to generate millions of transformants in E. coli EC100D. We termed this barcoded plasmid library pAD280_NN1 (strain AMD3507 in the EC100D background). We verified that the plasmid was fully barcoded by sequencing the plasmids from 10 random clones and found that all contained a single unique DNA barcode. To estimate the true barcode diversity of pAD280_NN1, we performed BarSeq on the plasmid library. Using an alpha diversity statistic, we estimate the true diversity of the pAD280_NN1 library is >10 million. To enable delivery of pAD280_NN1 by conjugation, we transformed the pAD280_NN1 library into electrocompetent E. coli WM3064 cells and selected for transformants in LB supplemented with carbenicillin (50 µg/mL) and DAP (400 µM), resulting in strain library AMD3510.

We generated an RB-TnSeq mutant library in PA14 via conjugation with AMD3510. To prepare the cells for conjugation, PA14 was grown across 5 LB agar plates at 37°C for 16 hours, while an aliquot of AMD3510 was grown in 50 mL of LB supplemented with carbenicillin (50 µg/mL) and DAP (400 µM) at 37°C for 6 hours until the cells reached mid-log phase. We scraped the PA14 cells from the 5 LB plates into 30 mL of LB + DAP, and after washing the AMD3510 cells to remove the residual carbenicillin, we combined donor cells with the PA14 cells to a final ratio of 1:1. We then conjugated this mixture on 10 separate LB + DAP plates for 5 hours at room temperature. We then scraped this conjugation mixture into 20 mL of LB + 15% glycerol, made 20 separate 1-mL aliquots in cryovial tubes, and then stored these at -80°C. To gauge the mutagenesis efficiency, we subsequently thawed one conjugation aliquot, did a dilution series, plated these on LB plates supplemented with 30 µg/mL gentamicin, and grew the plates at 30°C. The following day we counted colonies and calculated how many conjugation aliquots and LB + gentamicin plates we would need to make a diverse RB-TnSeq library. For the final scale-up, we pooled colonies from 120 LB + gentamicin plates (each plate contained a few thousand colonies) into 20 mL of LB + gentamicin. We then back-diluted this mixture into 150 mL of fresh LB + gentamicin to a starting OD of 0.75 and grew the culture at 30°C for 3 hours to allow the population to divide one time. We added glycerol to a final concentration of 15%, made dozens of single-use aliquots in cryovial tubes (2 mL per aliquot), and collected cell pellets for gDNA extraction. To map the transposon insertion locations and link these to their associated DNA barcodes, we followed a two-step approach to enrich transposon junctions (73). To increase the coverage of our mapping, we performed two independent library preps and sequenced these on an Illumina NovaSeq instrument (Novogene). The final *P. aeruginosa* PA14 RB-TnSeq mutant library (Paeruginosa_PA14_ML18; AMD3559) contains 446,458 uniquely barcoded and confidently mapped insertions (30). For genome-wide fitness assays, we performed DNA barcode sequencing (BarSeq) as described previously (30, 31), except we used dual-indexed primers to account for instances of index hopping on the Illumina NovaSeq instrument.

All software and analysis methods were performed as described in (30) and can be found in https://bitbucket.org/berkeleylab/feba/src/master/.

### Survival assay in large electrochemical bioreactors

The survival assay was performed with minimal adjustments from what was reported in (16, 17, 26). Briefly, a 1 mL mid-exponential (OD_500_ = 0.5) frozen aliquot of the RB-TnSeq library was added to a large LB 250 mL culture and grown at 30°C for ∼12 hours until a final OD_500_ = 3.0 ± 0.25. These cells were pelleted (6800 x g, 8 minutes) and washed twice in our MOPS minimal medium (100 mM MOPS pH 7.2 with NaOH, 43 mM NaCl, 93 mM NH_4_Cl, 3.7 mM KH_2_PO_4_, 1 mM MgSO_4_). Finally, they were resuspended to a final OD_500_ = 75 and 1 mL was transferred to an mBraun glovebox (Unilab model, N_2_ only) with the main chamber of the prepared glass reactors containing 99 mL of MOPS minimal medium with 20 mM D-glucose, 3.6 μM FeSO_4_, and 75 μM PCN (ChemScene) (in given samples), sparged for 30 minutes with N_2_. A graphite working electrode (Alfa Aesar #14738), platinum mesh counter electrode (custom-made using Alfa Aesar platinum gauze #10283), and Ag/AgCl reference electrode (Basi #MW-2030) were used to connect the sample to a potentiostat (Admiral Instruments, Squidstat Prime). The counter electrode was kept in a separate side chamber separated from the rest of the culture by a glass frit that contained 9 mL minimal medium. The cultures were stirred at 150 rpm and maintained at a temperature of 33°C (JOANLAB MHS4Pro) over the course of the experiment. The working electrode was poised at a potential of 0 mV v. the reference electrode. On sampling days, 100 µL was taken for CFU measurements and 1 mL was taken for outgrowth in LB. Outgrowth was performed by diluting 1 mL of culture into 20 mL LB and grown in LB at 37°C for 2 hours (∼4 generations). For gDNA preparations for sequencing, the Molecular Designs Column-Pure Bacterial Genomic DNA Kit (D423-100) was used.

### Survival assay in 96-well potentiostat system

7 mL, 20 hour LB cultures were pelleted and washed twice in MOPS minimal medium. Cells were resuspended to an OD_500_ of 75 and transferred into an anoxic Coy Chamber (33°C, 2-3% H_2_) containing the 96-well electrochemical plate containing a carbon working electrode/counter electrode and Ag/AgCl reference electrode (26, 74). The cultures were diluted 1:5 in N_2_-sparged minimal medium and 10 µL was inoculated into a well containing 190 µL glucose minimal medium. Both a slit-seal cover (BioChromato #R80.120.00) and an aluminum seal cover (DiversifiedBiotech #ALUM-1000) were added to the top of the plate to avoid evaporation during the experiment as well as maintain sterility. The plate was kept at 33°C with no shaking in the custom 96-well potentiostat system. Every well’s potential was maintained at 0 mV v. Ag/AgCl. On sampling days, the aluminum seal was removed and a pipetting volume of 100 µL was used to mix the contents of the plate. 10 µL of sample was removed for CFU measurements. The plate was resealed with a fresh aluminum seal and placed back into the 96-well potentiostat system to continue the assay.

### ATP measurement assay

The assay was performed as described in (17) with slight adjustments. Briefly, 20 µL of culture was removed from the potentiostat system on day 1 of the experiment and added to 180 µL of DMSO to quench/dissolve the cells. The sample was further diluted into 800 µL of 100 mM HEPES (pH = 7.5). 25 µL of the sample was mixed with 25 µL of the Promega BacTiter-Glo reagent in a 96-well black microtiter plate. The plate was incubated for 15 min at 30°C and luminescence was measured using a Tecan Spark 10M microplate reader. A standard curve derived from known concentrations of ATP was used to determine sample ATP concentrations.

### Data analysis

To compare fitness determinants across conditions, per-gene fitness scores from our PCN-cycling dataset were integrated with published TnSeq data from Basta *et al.* (28), who reported per-replicate read ratios across two replicates per condition for O_2_-lim and C-lim. Read ratios from Basta *et al.* were converted to log_2_ space by applying log_2_(read ratio + 0.001), where the pseudo-count of 0.001 was chosen to be at least 2-fold below the minimum observed nonzero ratio in either condition, minimizing distortion of real fitness values while assigning complete-dropout loci a defined limit. For each condition, gene-wise significance was assessed using a moderated two-sided t-test (75), which stabilizes gene-wise variance estimates by borrowing information across all genes, which was particularly important given the small number of replicates (n = 3 for PCN-cycling, n = 2 for O_2_-lim and C-lim). Fitness scores for each gene were calculated as the median of replicate values within each condition. Datasets were merged on PA14 locus tags after removal of intergenic regions from the Basta *et al.* dataset, yielding 4634 genes measured in all three conditions. Genes were classified as fitness-relevant if they met thresholds of fitness < −0.6 and p < 0.05 in a given condition. Overlap between conditions was computed in R and visualized as an Euler diagram using the *eulerr* package (v7.1.0).

To calculate cell-specific metabolic rate, current between days 1-7 was integrated (excluding the first 24 hours of the experiment as well as the 3 hours immediately following plate removal for CFU sampling to exclude current spikes immediately after returning the plate to the potentiostat), producing values with units nA*hr. This was converted to electrons using Faraday’s constant and divided by the geometric mean of the CFU measurements from days 1,3,5,7, multiplied by the well volume of 200 µL as well as total hours the experiment was run for. This led to a measurement of cell-specific metabolic rate in units of electrons sec^-1^ cell^-1^, as was calculated/reported in (26).

To calculate integrated charge over the first 24 h of anaerobic survival, current during this window was integrated to produce values with units of nA·hr, using the same current-integration approach described above for CSMR but restricted to the 0–24 h window. Unlike CSMR, integrated charge over the first 24 h is reported directly in nA·hr without normalization to CFU counts.

All statistical testing as well as plotting was performed in R 4.5.1, and the limma (75) and *eulerr* (v7.1.0) packages were used for the analysis presented in Fig. 2. Claude Sonnet 4.6 was used solely to generate code for appropriate statistical testing and figure generation. Geometric mean was used as a summary statistic for metabolic rate measurements to account for the counting of CFUs in log space. Replicates of a condition were linearly averaged otherwise.

## Acknowledgements

We thank Dr. Megan Bergkessel for generating the Δ*rnr* strain, Dr. Inês B. Trindade, Dr. Xiaoyu Shan, and Dr. Rob Phillips for help with data analysis and experimental design, and members of the Newman Lab for thoughtful discussions and feedback.

## Author Contributions

K.D. and D.K.N. conceived and designed the study. K.D. performed the experiments and analyzed the data. R.A.M. and H.K.C. performed sequencing and analysis of RB-TnSeq data. A.M.D. provided resources and expertise for RB-TnSeq library construction and BarSeq methodology. A.O. provided resources and expertise for the electrochemical setup. K.D. and D.K.N. wrote the manuscript with input from all authors. D.K.N. supervised the project and secured funding.

---

**Supplemental Fig. 1.**
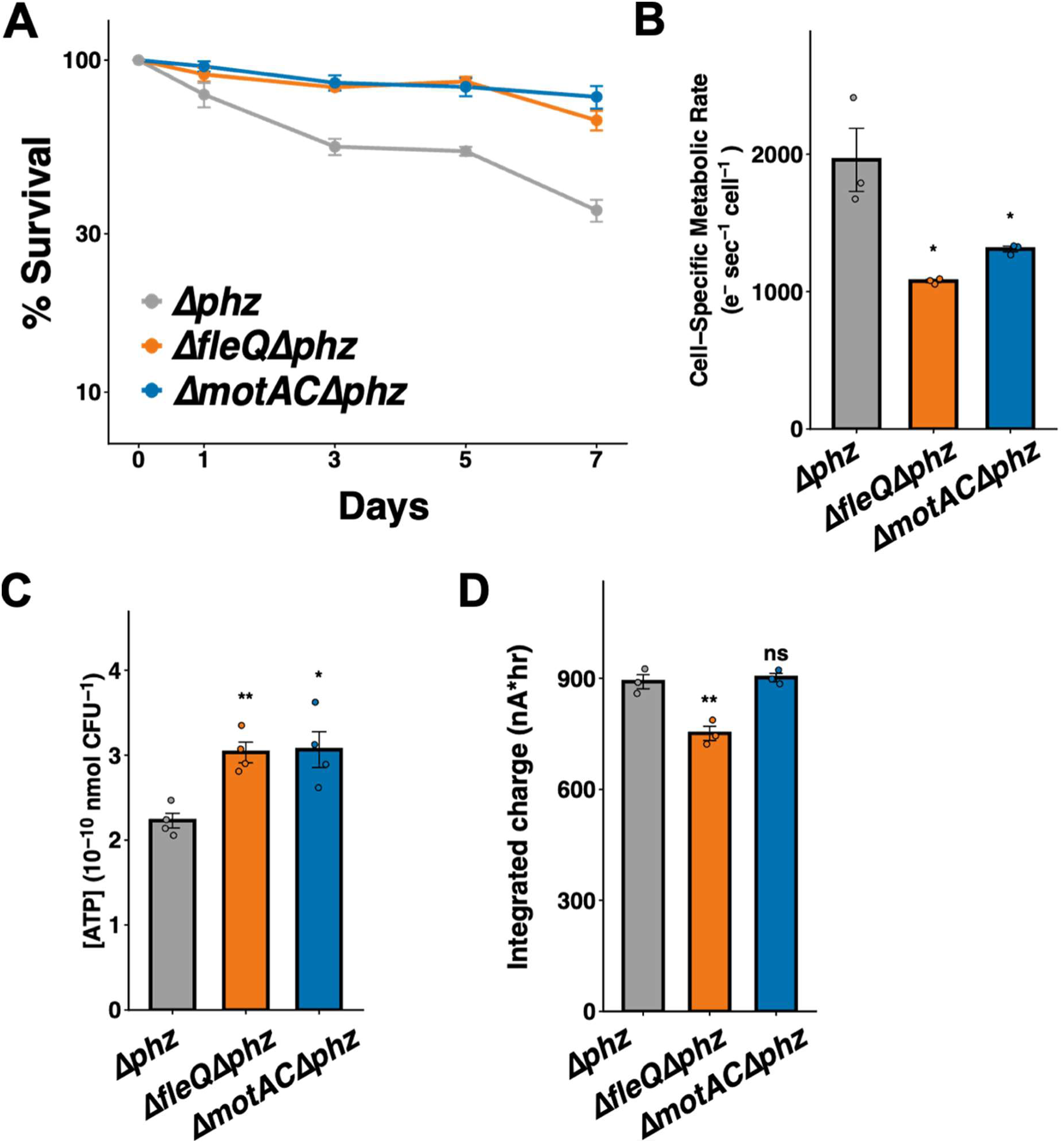
Loss of MotAC improves survival, metabolic rate, and ATP levels in the *Δphz* background. **(A)** Survival of *Δphz*, *ΔfleQΔphz*, and *ΔmotACΔphz* strains over seven days of anaerobic PCN-cycling survival, expressed as percent survival relative to day 0. Data points represent the mean of n = 3 biological replicates; error bars indicate standard error of the mean. **(B)** Cell-specific metabolic rate (CSMR, electrons·cell⁻¹·sec⁻¹) of the indicated strains measured during anaerobic PCN-cycling survival. **(C)** Per-cell ATP concentrations (nmol CFU⁻¹) of the indicated strains on day 1 of anaerobic survival. **(D)** Integrated charge (nA·hr) over the first 24 h of anaerobic survival. For panels B and D bars represent the mean of n = 3 biological replicates while bars in panel C represent the mean of n = 4 biological replicates; error bars indicate SEM; individual data points are shown. Statistical comparisons are relative to *Δphz*; *, p < 0.05; **, p < 0.01.

